# Bilingual speakers’ enhanced monitoring can slow them down

**DOI:** 10.1101/2021.05.12.443807

**Authors:** Roksana Markiewicz, Ali Mazaheri, Andrea Krott

**Author notes:** **Data availability:** Stimuli and data are available on request. **Authorship Contributions:** AK conceptualised the study. RM collected the data and analysed the data. AK and AM supervised data analyses. RM wrote the manuscript. AK supported the manuscript writing. All authors edited the manuscript.

## Abstract

Performance differences between bilinguals and monolinguals on conflict tasks can be affected by the balance of various sub-processes such as conflict monitoring and allocation of attentional resources for stimulus categorisation. Here we investigated the effect of bilingualism on these sub-processes during a conflict task with medium monitoring demand. We examined the behavioural responses and evoked potentials from bilinguals and monolinguals during a flanker task with 25% incongruent trials. We analysed behavioural differences by means of averaged response times and ex-Gaussian analyses of response time distributions. For the evoked potentials we focused on N2 (reflecting conflict monitoring) and P3 responses (reflecting allocation of attentional resources for cognitive control). We found that bilinguals had significantly longer response distribution tails compared to monolinguals. Additionally, bilinguals exhibited a more pronounced N2 and smaller P3 components compared to monolinguals, independent of experimental condition, suggesting a different balance of sub-processes for the two groups. It is suggested that bilinguals engaged more strongly in monitoring processes, leading to the allocation of fewer attentional resources during stimulus categorisation. Importantly, N2 amplitudes were positively and P3 amplitudes negatively related to the length of response distribution tails. We postulate that these results reflect an overactive monitoring system in bilinguals. This enhanced monitoring led to reduced engagement of attentional resources for stimulus categorisation, but also occasionally to slow responses. These results suggest that changes of the cognitive control system due to bilingual experience can change the balance of processes during conflict tasks, potentially leading to a small behavioural disadvantage.

## Introduction

Speaking another language leads to changes in brain structure and function, particularly in the networks implicated in cognitive control (Li et al., 2014; Pliatsikas & Luk, 2016). Consequently, differences have been found between monolingual and bilingual speakers in tasks of executive control, with many studies finding superior behavioural performance for bilinguals (Bialystok et al., 2004; Costa et al., 2008; Emmorey et al., 2008; Hernández et al., 2010; review in Bialystok et al., 2012; meta-analyses in Donnelly et al., 2019; Grundy, 2020).

However, other studies have found similar performance in the two participant groups or even superior performance in monolinguals (Duñabeitia et al., 2014; Gathercole et al., 2014; Naeem et al., 2018; Paap et al., 2015; review in Nichols et al., 2020; meta-analysis in Lehtonen et al., 2018). Together these findings suggest that structural and functional changes brought about by bilingualism lead to differences in cognitive processing, but this is not always reflected in a behavioural advantage. Importantly, cognitive control tasks contain a number of processes. Functional and structural changes might affect these processes in different ways. They might make bilinguals more efficient in some and potentially less efficient in other processes involved in a task. The current study investigated the differences in engagement of sub-processes of cognitive control between monolingual and bilingual speakers and examined how these could lead to behavioural differences.

One feature of executive function tasks that has to date not much been considered as a potential source of inconsistencies in the literature is that they always involve more than one process, such as conflict monitoring or engagement of attentional resources for conflict control. If bilinguals are more engaged in some processes and less engaged in others, then the balance of engagement of these processes in a particular task could affect behavioural differences between bilinguals and monolinguals. If a process in which bilinguals are more engaged in is beneficial for performance in a particular task, then bilinguals will outperform monolinguals. If a process in which bilinguals are more engaged in is balanced with a process in which they are less engaged in, the cost and benefits of the processes might cancel each other out and there will be no performance difference between the participant groups. And finally, if a process is beneficial for performance in which bilinguals are less engaged in, then monolinguals might outperform bilinguals. The balance of processes involved in a task might therefore potentially explain inconsistencies in the literature.

One very common cognitive control task, and one that we used in the present study, is the flanker task (Eriksen & Eriksen, 1974). In the arrow version of this task, participants are presented with rows of arrows and need to indicate by button presses whether the central arrow (the target) points to the left or right. The flanking arrows are either congruent with the central arrow (pointing into the same direction as the target arrow) or incongruent (pointing into the opposite direction of the target arrow). The incongruent condition creates a conflict that needs to be resolved. It therefore leads to longer response times and more errors than the congruent condition.

Much of the literature on the flanker task has studied the processes that lead to more errors and slower responses for incongruent than congruent flankers (see overview in Ridderinkhof et al., 2021). In their seminal study, Eriksen and Eriksen (1974) suggested that both target and flankers are processed in parallel and activate corresponding responses, leading to response competition (see ‘continuous flow’ model by Eriksen & Schultz, 1979). Based on electrophysiological measures, Gratton and colleagues (Gratton et al., 1988, 1992; Coles et al., 1985) developed this idea into a two-phase model of stimulus evaluation: during an early phase (150-250ms) (“parallel phase”), both flankers and the target affect the response. In the later phase (“focused phase”), participants focus on the location of the target and their responses are strongly affected by the target. Ridderinkhof et al. (1995) suggested a dual-route model. Their ‘direct’ route is very similar to the parallel phase of the two-phase model. But target selection and identification as well as stimulus-response translation is handled by a slower ‘deliberate’ route. These two routes are activated in parallel and both prime a response. Taken together, all models assume some form of merging and competition of responses. The resolution of the competition requires cognitive control mechanisms. Since the models are relatively similar and can account for most phenomena in the literature, we will restrict our discussion in what follows to the more recent dual-route model.

Important for the present study, processing in the flanker task is under strategic control (Gratton et al., 1992). Participants may rely more heavily on information from the direct route versus the deliberate route (see dual-route model). This suggestion is based on the finding that participants process the same stimuli differently in different situations. For instance, incongruent flankers interfere more strongly when congruent trials are dominant in a list (75%) than when congruent and incongruent trials occur equally often (50/50) (e.g., Gratton et al., 1992). Such strategic effects have been attributed to the modulation of conflict monitoring (Botvinick et al., 2004). When trials are predominantly congruent, participants rely more strongly on the direct route and monitor less for conflict. This leads to faster responses for congruent trials. But due to reduced conflict monitoring, processing of incongruent trials gets more erroneous, effortful and slower. In other words, a conflict monitoring mechanism detects conflict and upregulates cognitive control to resolve the conflict. This explains how participants can upregulate control when needed (Bugg & Gonthier, 2020), for instance as a proactive strategy in predominantly incongruent lists. The fact that conflict monitoring is not consistently used suggests that monitoring is effortful and is avoided when not necessary.

Some studies have found superior performance of bilingual compared to monolingual adults on the flanker task (Abutalebi et al., 2015; Costa et al., 2008, 2009; Emmorey et al., 2008; Zhou & Krott, 2016). But the mixture of congruent and incongruent trials makes a difference (Costa et al., 2009). Superior bilingual performance seems to only occur when incongruent trials occur with a high (50%) or medium (25%) probability, not when one trial type (congruent or incongruent) dominates (92%). In terms of the sub-processes of the flanker task, this suggests that bilinguals might follow different strategies depending on the mixture of the trial types. When one type of trials dominate, bilinguals seem to follow the same strategy as monolinguals. If congruent trials dominate, both groups likely rely more heavily on the direct route of processing. If incongruent trials dominate, both groups likely rely more heavily on the deliberate route that involves cognitive control of the competing responses. Group differences occur for lists of 50% or 25% incongruent flankers. In this case, it is presumably most advantageous to assess whether the stimulus contains a conflict or not so that the deliberate route can be given priority and cognitive control processes can be engaged in case of conflict. In other words, upregulated conflict monitoring can lead to better performance in the incongruent condition. Therefore, bilinguals seem to outperform monolinguals in terms of conflict monitoring, but only when conflict monitoring can make a difference to task performance. Bilinguals might possess a more efficient monitoring system or engage their monitoring system more strongly than monolinguals. Monolinguals, in contrast, might avoid constant monitoring more strongly because it is too effortful.

But not all studies have reported behavioural differences between bilingual and monolingual adults in the flanker task (Antón et al., 2019; Grundy, Chung-Fat-Yim, et al., 2017; Kousaie & Phillips, 2012, 2017; Luk et al., 2010). While it is not possible to pinpoint one particular difference that explains the inconsistencies across studies, differences in conflict monitoring demands across studies might be one of the reasons. For instance, Grundy, Chung-Fat-Yim, et al. (2017) presented only one flanker on each side of the target. It might be easier to focus on the target in this case and a more engaged monitoring mechanism for bilinguals might not be beneficial. Antón et al. (2019) presented only 16 trials per condition. Due to response variability, group differences might not be present. They also presented the flanker task as part of a series of tasks (see also Kousaie & Phillips, 2012; 2017), which might alter conflict monitoring in case of carry-over effects (see, e.g., carry-over effects on stop-signal task performance in Kałamała et al., 2022). And Kousaie and Phillips (2012; 2017) mixed congruent and incongruent trials with neutral trials (targets without flankers), which again might have altered the engagement of conflict monitoring because conflict monitoring was only needed for trials with flankers.

It is common practice to assess performance on tasks investigating cognition, such as the flanker task, through averaged response times. This method has the disadvantage, though, that it cannot assess the involvement of different processes (e.g., automatic versus controlled processes) involved in the task. One approach that partly addresses this limitation is to examine response time distributions instead of averaged response times. This has been done in the bilingual advantage literature with the means of ex-Gaussian analyses (Abutalebi et al., 2015; Calabria et al., 2011; Tse & Altarriba, 2012; Zhou & Krott, 2016). By utilising an ex-Gaussian analysis, it is possible to separate a measure of general processing speed from a measure of slow responses for a particular participant in a particular experimental condition. General processing speed is captured by the μ parameter (the mean of the Gaussian part of the response distribution), the variance of the Gaussian part is reflected by σ, and the extremeness and frequency of slow responses is reflected by *τ* (the mean and variance of the exponential part of the response distribution). While ex-Gaussian parameters do not reflect specific processes, μ is commonly interpreted as reflecting more automatic processes, while *τ* is interpreted to reflect controlled processes that are attention demanding (see review in Matzke & Wagenmakers, 2009). This fits the distinction made in the flanker literature between responses that more heavily rely on the direct route of processing on the one hand and responses that more strongly rely on the deliberate route of processing on the other hand. This means that μ is more strongly affected by the direct processing route of the stimuli in the flanker task and τ is strongly affected by the deliberate processing route. But one needs to be careful not to over-interpret μ because it is still a reflection of general processing speed, which includes both responses based more strongly on automatic processes and responses based more strongly on controlled processes. Also, since the flanker task involves various controlled processes, τ might not only be affected by the deliberate processing route, but also by conflict monitoring or the control processes needed to merge the outcome of the deliberate route with that of the direct route, which often involves inhibitory control processes.

Investigations into ex-Gaussian parameters for conflict tasks have found larger μ in conflict conditions compared to no-conflict conditions (e.g. Heathcote et al., 1991; Kane & Engle, 2003; Spieler et al., 2000; Zhou & Krott, 2018). This has been interpreted as a reflection of differences in response conflict and inhibitory control processes across the two conditions (Heathcote et al., 1991; Kane & Engle, 2003; Spieler et al., 2000; Zhou & Krott, 2018). In contrast, *τ* is generally not affected by conflicting information. It is often thought to reflect attentional control (Epstein et al., 2011; Hervey et al., 2006; Spieler et al., 1996). Given the sub-processes of the flanker task, we can be more concrete about how attentional control can affect τ in this task. As indicated, particularly slow responses likely occur when participants rely strongly on controlled stimulus processing. This could be an over-reliance on conflict monitoring, meaning a response might be slowed down due to overactive monitoring processes. It could also be that a controlled process is not very efficient. For instance, fewer attentional resources might be devoted to the cognitive control mechanisms for the deliberate processing route or the merging of direct and deliberate processing routes of the flanker task.

Utilizing ex-Gaussian analyses to investigate effects of bilingualism in conflict tasks has suggested differences between language groups, particularly in terms of controlled processes. Bilingual speakers have been found to have smaller μ and *τ* (Abutalebi et al., 2015; Calabria et al., 2011) or only smaller *τ* (Zhou & Krott, 2018). Smaller *τ* (shorter response distribution tails) for bilinguals has been interpreted as an advantage in attentional control. As indicated above, this could mean bilinguals engage in more efficient conflict monitoring. Alternatively or in addition, they might have more efficient cognitive control processes related to the deliberate processing route or the merging of the two processing routes of the conflict task.

Differences in functional processing between monolingual and bilingual participants are rather difficult to determine merely on the basis of behavioural results, even when employing ex-Gaussian analyses. Brain imaging provides better methods for this purpose. Previous EEG research into the bilingual advantage in conflict tasks has found differences particularly in the N2 and P3 components. The N2 component is a fronto-central negative deflection that peaks at around 200-300 ms after stimulus onset (Grundy, Anderson, et al., 2017). Its neural source has been localised to the anterior cingulate cortex (ACC) (e.g., Yeung et al., 2004). The ACC is an integral part of the cognitive control system and has been associated with conflict monitoring (Botvinick et al., 2004; Carter et al., 1998). It has been suggested that the ACC detects conflict and engages the dorsolateral prefrontal cortex to resolve the conflict (Carter & van Veen, 2007). Because of this and experimental findings, the N2 is associated with conflict monitoring and detection. Conditions that involve conflict, for instance incongruent trials in the flanker task, elicit a more prominent N2 response compared to conditions that are conflict-free (i.e. congruent trials) (Danielmeier et al., 2009; Purmann et al., 2011; van Veen & Carter, 2002; Wild-Wall et al., 2008; Yeung et al., 2004). The amplitude of the N2 is sensitive to the monitoring demand. N2 amplitudes are decreased for flankers that are more distant to the target in the flanker task (Danielmeier et al., 2009) and the congruency effect on N2 is dependent on the frequency of the incongruent trials in the flanker task. It only occurs when incongruent trials are infrequent (e.g., 25%), not when they are frequent (75%) (Purmann et al., 2011), thus during heightened monitoring demand. The N2 also shows conflict adaptation effects, with smaller N2 amplitudes for incongruent trials that follow incongruent trials compared to congruent trials (e.g., Clayson & Larson, 2011; Forster et al., 2011)(Clayson & Larson, 2011; Forster et al., 2011). A larger N2 amplitude is thus associated with enhanced conflict monitoring (Folstein & Van Petten, 2008; Kousaie & Phillips, 2012; Yeung & Cohen, 2006). More concretely, given the association of the N2 with the ACC and the role of the ACC in the conflict control loop to signal the dorsolateral prefrontal cortex to increase cognitive control (e.g., Carter & van Veen, 2007), the N2 has been suggested to reflect this signalling for cognitive control (Clayson & Larson, 2011).

Bilinguals have often been found to show more pronounced or earlier N2 responses compared to monolinguals (Chung-Fat-Yim et al., 2021; Fernandez et al., 2013, 2014; Kousaie & Phillips, 2012, 2017; Morales et al., 2015; Moreno et al., 2014). More pronounced N2 components have been found particularly in Go/No Go and AX-CPT tasks (Fernandez et al., 2013, 2014; Morales et al., 2015; Moreno et al., 2014). N2 effects in bilinguals have also been found to be moderated by L2 proficiency, with higher L2 proficiency leading to more pronounced N2 amplitudes (Fernandez et al., 2013, 2014). In line with the interpretation of the N2, larger N2 amplitudes in bilinguals echo greater resource allocation to conflict monitoring and shorter N2 latencies suggest more automatic and faster monitoring. Both differences suggest that bilinguals rely more strongly on earlier processes during a conflict task than monolinguals, reflecting a more proactive approach (Grundy, Anderson, et al., 2017). In contrast to the former studies, Kousaie and Phillips (2012, 2017) found more pronounced or earlier N2 amplitudes in monolinguals compared to bilinguals, the former for young adults in a Stroop task, the latter for older adults in a Simon task. The Stroop result could be due to the verbal nature of the task and therefore the diversion of resources to linguistic processes in bilinguals. But as the authors interpreted the results, smaller N2 amplitudes in bilinguals could mean less requirement for conflict monitoring in bilinguals to perform the task. Importantly, as mentioned above, the exact task features can affect group differences. It is therefore maybe not that surprising that different tasks and variations of tasks have led to different results.

The other ERP component relevant to the present study is the P3 component (also called standard P3 or P3b), a positive component with centro-parietal distribution that is thought to be the summation of activity widely distributed in the brain (Johnson, 1993). The P3 is typically elicited during processes of decision making and associated with attentional resource allocation during stimulus categorisation. Shorter P3 latencies are associated with shorter stimulus evaluation / faster categorisation time (Coles et al., 1985; Kok, 2001; Polich, 2007), and larger amplitudes are proportional to the amount of attentional resources allocated to stimulus processing (Kok, 2001; Wickens et al., 1983) or, more specifically for the flanker task, to the amount of attentional resources needed for cognitive control during stimulus categorisation (Clayson & Larson, 2011). Incongruent trials in the flanker task lead to larger and later P3 responses (Purmann et al., 2011; Wild-Wall et al., 2008). But, similar to the N2, this difference is dependent on the demand for control. An amplitude difference only occurs when incongruent trials are infrequent (e.g., 25%), not when they are frequent (75%), and a peak latency difference is larger in the infrequent condition (Purmann et al., 2011). Also, the P3 in the flanker task increases after conflict trials and is therefore sensitive to the recruitment of attentional resources needed for improved cognitive control during stimulus categorisation (Clayson & Larson, 2011).

Again, differences in P3 between bilinguals and monolinguals seem to depend on the task. Bilinguals have been found to show greater P3 amplitudes (Kousaie & Phillips, 2017; Moreno et al., 2014) and/or shorter P3 latencies than monolinguals in conflict tasks (Barac et al., 2016; Kousaie & Phillips, 2012, 2017). In contrast, other studies showed the opposite language group effect, with bilinguals producing smaller P3 amplitudes during a Stroop task (Coderre et al., 2014), a Simon task (Kousaie & Phillips, 2012), and a Flanker task (Botezatu et al., 2021). These findings taken together suggest whether bilinguals show a larger or smaller P3 response and therefore more or less activation of attentional resources for cognitive control during stimulus categorisation depends on the task.

The task dependency of changes to cognitive control in bilingual speakers is also evident in studies that manipulated language context during a conflict task. Wu and Thierry (2013) tested bilingual speakers on a flanker task in which flanker trials were interleaved with words from either one or both of the speakers’ languages. They found reduced P3 amplitudes in the mixed language condition compared to the single language conditions (but see Chung-Fat-Yim et al., 2021). This effect was replicated by Jiao et al. (2020) who replaced the interleaved word comprehension task with an interleaved picture naming task. This study found an additional increase in terms of the N2 component in the mixed condition. In a related study, P3 amplitudes have been found to be reduced in a stop-signal task after bilinguals used their L2 (Kałamała et al., 2022), that is after enhanced engagement of inhibitory control processes. These results suggest that conditions of enhanced control demand increase bilinguals’ N2 components and reduce their P3 responses. Increased N2 amplitudes suggest a stronger engagement of conflict monitoring processes, while reduced P3 amplitudes suggest the devotion of fewer attentional resources for cognitive control during stimulus processing, thus more efficient categorisation.

## The present study

The aim of the present study was to investigate differences in engagement of sub-processes of cognitive control between monolingual and bilingual speakers and how these might lead to behavioural differences. For that, we tested young adults on a flanker task. In order to detect group differences in more automatic versus controlled processes, we inspected ex-Gaussian parameters. But more importantly, we investigated functional neural differences between the two participant groups by studying event-related potentials during task performance, focussing on N2 and P3.

Costa et al. (2009) found a bilingual advantage on a flanker task only when monitoring demand was medium or high. In the current study, we investigated group differences in a version with medium monitoring demand; that is with 25% incongruent and 75% congruent trials. This version allows strategic flexibility and is therefore particularly suitable for finding group differences in terms of conflict monitoring or of engagement of attentional resources for cognitive control. Given previous results, we expected to see stronger bilingual monitoring reflected in enhanced N2 amplitudes. Jiao et al. (2020) showed that stronger engagement of monitoring processes in bilinguals in a flanker task, that is increased N2 responses, went hand in hand with decreased P3 amplitudes. We therefore might also see decreased P3 amplitudes in bilinguals and thus less subsequent devotion of attentional resources for cognitive control during response selection. We did not have strong predictions about behavioural results. If bilinguals are, as previously suggested, particularly efficient in monitoring in the task, they might outperform monolinguals in terms of accuracy and RTs and they might have shorter response distribution tails. However, with a relatively small percentage of incongruent trials (25%), strong bilingual engagement in monitoring might not be beneficial. In this case, we might not see group differences (Antón et al., 2019; Grundy, Chung-Fat-Yim, et al., 2017; Kousaie & Phillips, 2012, 2017; Luk et al., 2010) or even a monolingual advantage.

In order to investigate whether stronger or weaker engagement of sub-processes in the task is beneficial for performance or not and whether this is different for the two participant groups, we analysed whether neural markers that showed differences between groups correlate with ex-Gaussian parameters and whether these correlations differed between groups. Yeung and Nieuwenhuis (2009) had found that larger N2 amplitudes in a flanker task were correlated with slower responses. This means that a strong engagement of monitoring processes in bilinguals, reflected in larger N2 amplitudes, could potentially slow down their responses compared to monolinguals, meaning that a strong engagement of monitoring might be disadvantageous (see above). On the other hand, the need for fewer attentional resources for cognitive control during stimulus categorisation, reflected in decreased P3 amplitudes, might counteract this disadvantage. Since controlled processes such as conflict monitoring or the allocation of attentional resources for conflict control are rather reflected in the τ parameter, we might see that N2 and P3 are correlated with τ but not μ.

## Method

### Participants

Seventy students of the University of Birmingham took part in the study. Participants were allocated to one of two language groups (monolinguals/bilinguals) (see classification criteria in Table 1) based on their responses to a Language History questionnaire (taken from Zhou & Krott, 2018). 16 participants were excluded from the analysis due to: (a) EEG software malfunction (*N=7*), (b) excessive EEG artifacts (*N=5*, rejected trials exceeded 30%), or (c) extremely slow responses (*N=4*, RT mean was 3SD above the mean of all participants). This led to 28 monolinguals and 26 bilinguals being included into analyses (see Table 2 for demographic information of participants). All had normal or corrected vision and no neurological impairments. Monolinguals matched their bilingual counterparts in age, *t*(52) =.59, p>.05, and fluid intelligence, assessed via Raven’s Standard Progressive Matrices (Raven, 1958), *t*(52) =.56, *p*>.05.

**Table 1.** Allocation criteria into monolingual and bilingual groups.

| Monolinguals | Bilinguals |
| --- | --- |
| a) No second language before age 7 | a) Both L1 and L2 learnt before age 7 |
| b) Language proficiency rating 4 or lower (out of 7) in L2 (or any other language) | b) Language proficiency rating above 5 (out of 7) in both L1 and L2 |
| c) Use of only L1 on a daily basis | c) Both L1 and L2 used on a daily basis |

**Table 2.** Demographic information of participants.

|  | Monolinguals | Bilinguals |
| --- | --- | --- |
| N (Male/Female) | 28 (10/18) | 26 (3/23) |
| Mean Age in years (*range, SD*) | 20.1 (18-24, 1.66) | 19.8 (18-28, 2.04) |
| Right handed/ left handed | 24/4 | 26/0 |
| Mean age (years) of second language acquisition (*SD*) | 10.14 (2.91) | 3.54 (2.8) |
| Self-rated proficiency of L1 (*SD*) (range 0-7) | 6.71 (0.42) | 6.54 (0.65) |
| Self-rated proficiency of L2 (*SD*) (range 0-7) | 2.29 (0.76) | 6.15 (0.92) |
| Mean Raven's score (*SD*) | 50.68 (5.12) | 49.25 (9.04) |

Notes. Handedness was measured using the Handedness questionnaire by Oldfield (1971).
Seven of the monolingual participants reported some knowledge of another language.

### Materials

The flanker paradigm (Eriksen & Eriksen, 1974) was implemented using E-prime 2.0 (Psychology Software Tools, Pittsburgh, PA). Each trial consisted of five black arrows presented in the centre of a white computer screen, with the central arrow being the target and two flanking arrows on both sides functioning as distractors. Two types of trials were presented: (a) congruent trials (75% of all trials) which comprised of five arrows facing into the same direction and (b) incongruent trials (25% of all trials) which consisted of a central target arrow pointing to the opposite direction than the flankers. The Language History questionnaire (taken from Zhou & Krott, 2018) asked participants to self-rate their proficiency of English and any other languages that they learnt or were able to speak. They were also asked to report the age of acquisition of English and any other languages and their current language use pattern (e.g., using mainly one or using both languages on a daily basis at home/ school/ social setting).

### Procedure

Participants first completed the Language History questionnaire followed by the Handedness questionnaire (Oldfield, 1971), before taking part in the EEG experiment. For the latter, they sat in an isolated room, approximately 55cm from the computer monitor. Each trial began with the presentation of a fixation cross in the centre of the screen for 800 ms, followed by a stimulus, which stayed on screen for 5000 ms or until response. Each trial was followed by a blank screen for 500 ms. Subjects indicated the direction of the target by pressing corresponding buttons on a Cedrus RB-834 response pad, which also recorded response times. The task comprised of 10 blocks of 96 trials, with an additional practice block (24 trials), taking approximately 35 minutes. Stimuli appeared randomly. Finally, participants completed the Standardised Progressive Matrices assessment (Raven, 1958) for which they were given a 25 minute time limit.

### EEG recording

Continuous electrophysiological recordings were obtained using a 128-channel BioSemi Active Two EEG system. BioSemi electrodes are active electrodes with sintered Ag-AgCI tips/pallets. The 5-10 system (Oostenveld & Praamstra, 2001) was used for electrode positioning. External electrodes placed on the mastoids were used as reference electrodes for offline processing. The signal was amplified with a bandpass of .16-128 Hz using a Biosemi Active 2 AD-box, sampled at 512 Hz, and recorded using ActiView version 7.06 software.

## Data analysis

### Behavioural analysis

We analysed response times (RTs), accuracy, and ex-Gaussian parameters (μ and τ). We examined group differences using traditional measures (i.e., averaged RTs and accuracy) in order to easily compare the performance of our monolingual and bilingual groups to previous findings (see Supplementary material and Supplementary Figure 1 for detailed results). The accuracy rate was the percentage of correct responses. Responses below 200 ms were treated as incorrect. RT and ex-Gaussian analyses were based on correct responses. Responses greater than 2SD from each participant’s mean were removed from the RT analysis (for a similar approach see, e.g., Paap & Greenberg, 2013; Zhou & Krott, 2018), but not from the ex-Gaussian analysis. Ex-Gaussian parameters μ and *τ* were determined for each participant for each condition (congruent and incongruent) using the QMPE software (Brown & Heathcote, 2003). The exit codes were below 32 for all parameter estimations, therefore they were considered trustworthy (in line with the QMPE software manual). The average number of search iterations was 7.09. RTs, accuracy, and ex-Gaussian parameters (μ and *τ*) were analysed each with a mixed 2 (Language Group: monolingual vs. bilingual) × 2 (Condition: congruent vs. incongruent) analysis of variance (see Suppl. Material for RT and accuracy results).

### EEG analysis

The EEG data pre-processing was performed with the means of EEGLAB 14.1.2b (Delorme & Makeig, 2004) and Fieldtrip toolbox 2018-07-16 (Oostenveld et al., 2011). EEG data were off-line filtered with a 0.1 Hz high pass filter and a 30 Hz low pass filter, referenced to the average of the mastoid electrodes and epoched from −2 to 2 seconds locked to the onset of the arrow array. Ocular artefacts in the data were removed based on the scalp distribution and time-course, using the independent component analysis (ICA) extended-algorithm in EEGLAB. Prior to the ICA, a Principal Component Analysis (PCA) was used to reduce dimensionality of the data to 15 components. The average number of removed components per participant was 1.33 (*SD* = .57, min = 1, max = 3), with no difference in the number of components removed between the bilingual (*M* = 1.5, *SD* = .65) and monolingual (*M =* 1.21, SD = .5) group (*t*(52) = −1.82, *p* > .05). The Fieldtrip toolbox 2018-07-16 (Oostenveld et al., 2011) was used for trial removal. Our criteria for trial rejection were: trials with incorrect responses, trials with RT < 200 ms, and trials with RT > 2SD from the mean of each participant. The average percentage of rejected trials per participant due to behavioural outliers (i.e., reasons outlined above) was 9.41% (SD = 2.39), with no significant difference in the % of trials removed between the bilingual (M=9.32, SD = 1.95) and monolingual (M = 9.5, SD = 2.77) groups (t(52) = .28, p >.05). In addition, a manual visual artifact rejection summary method was used to remove trials (per electrode) with a maximum value < 200 μV. The average percentage of rejected trials per participant due to artifact rejection was 7.82 (SD = 6.46), with no significant difference in the % of trials removed between the bilingual (M = 8.09, SD =6.75) and monolingual (M = 7.6, SD = 6.31), t(50) = −.28, p >.05). The total average percentage of rejected trials per participant included in the analysis was 16.0% (*SD = 6.6*), with no significant difference in the number of trials removed between the bilingual (*M = 15.96%, SD* = 6.8) and monolingual (*M = 16.1%, SD* = 6.45) groups (*t*(52) = .08, *p* > .05). The traditional RT and ERP analyses were conducted using (on average per participant) 620 (SD = 95.4) congruent trials and 177.5 (SD = 30.5) incongruent trials.

### Event-Related Potentials (ERPs)

We averaged the time-locked EEG activity of all valid trials (see above for removed trials) for congruent and incongruent conditions separately. The baseline correction used for the ERP analysis was −200 ms to 0 prior to stimulus (arrow array) onset. We employed a nonparametric cluster-based permutation test (Maris & Oostenveld, 2007) as implemented in Fieldtrip to assess statistical differences of ERP amplitudes in the time window 0-1 s after stimulus onset between: (1) conditions collapsed across all participants (congruent vs incongruent), (2) groups for each condition (monolinguals vs. bilinguals on congruent condition; monolinguals vs. bilinguals on incongruent condition), and (3) groups for flanker effect (i.e. congruency effect in monolinguals vs. congruency effect in bilinguals). Specifically this approach involved conducting a t-test (dependent samples in the condition contrast, independent samples in the group contrast) for every point in the electrode by time plane, and clustering the t-stats in adjacent spatiotemporal points if they exceeded a threshold of *p* < .05 (cluster-alpha). The cluster with a Monte Carlo *p*-value smaller than .025 was identified as significant (simulated by 1000 partitions), thus, showing a significant condition or group difference in amplitude. Probability values for each cluster were obtained by comparing it to a distribution using the Monte Carlo simulation method, in which the group (and equivalent condition) labels were randomly shuffled 1000 times and the maximum sum of the t-stat in the resulting clusters was measured. Here we considered clusters falling in the highest or lowest 2.5th percentile to be considered significant. The triangulation method was used to define a cluster (i.e. a cluster consisted of at least two significant neighbouring electrodes).

In order to investigate whether stronger or weaker engagement of sub-processes in the task was beneficial for performance or not and whether this was different for the two participant groups, we correlated the ex-Gaussian parameters μ and *τ* with markers of neural differences between groups (e.g. N2 and P3 amplitudes). In case of a significant correlation, we investigated whether the factor language group interacted with the neural marker in a regression analysis.

## Results

### Behavioural results

Figure 1 shows the distributions of average Ex-Gaussian parameters μ (panel A) and *τ* (panel B) of the flanker task for the two participant groups and the two conditions (see Supp. Fig. 1 for the distributions of average RT and accuracy).

**Figure 1.**
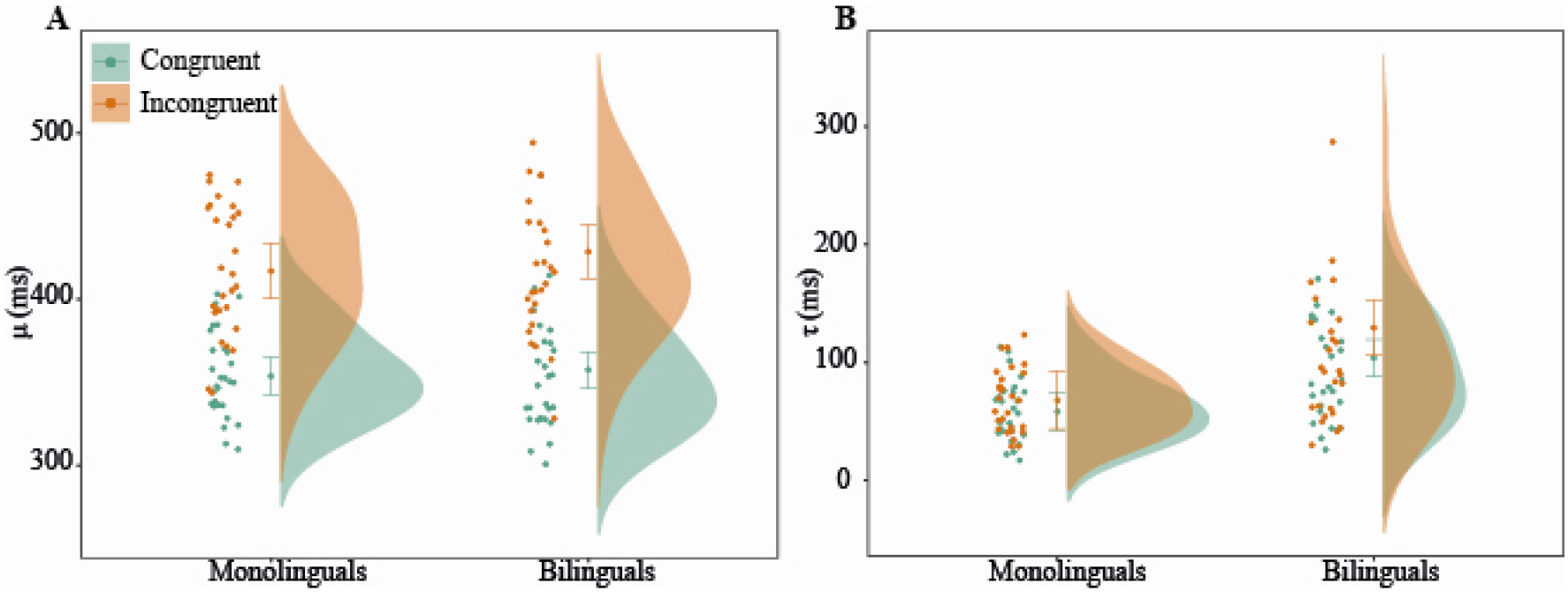
Distributions and means of Ex-Gaussian parameter μ (panel A), and Ex-Gaussian parameter *τ* (panel B) per condition (congruent and incongruent) in the flanker task for both monolinguals and bilinguals. Error bars represent 95% confidence intervals.

#### *Ex-Gaussian* μ

There was a significant main effect of Condition, F(1,52) = 441.35, p < .001, indicating larger μ in the incongruent compared to the congruent condition. There was no main effect of Language Group, *F*(1,52) = .69, *p =.411*, nor a Language Group by Condition interaction, *F*(1,52) = .1.53, *p = .222*.

#### Ex-Gaussian τ

There was a significant main effect of Condition, *F*(1,52) = 10.97, *p = .002*, indicating larger *τ* (i.e. longer distribution tails) in the incongruent compared to the congruent condition. There was also a significant main effect of Language Group, *F*(1,52) = 1.88, *p < .001*, with bilinguals having overall larger *τ* compared to monolinguals. The interaction between Condition and Language Group was not significant, *F*(1,52) = 2.36, *p = .13*.

### EEG results

#### ERP condition differences

Statistical analysis investigating ERP differences between congruent and incongruent trials revealed significant differences in the ERP waveform in time periods corresponding to the N2 and P3, as found previously (e.g., Bartholow et al., 2005; Purmann et al., 2011; Wild-Wall et al., 2008; Yeung et al., 2004), as well as a slow late component (see Figure 2).

**Figure 2.**
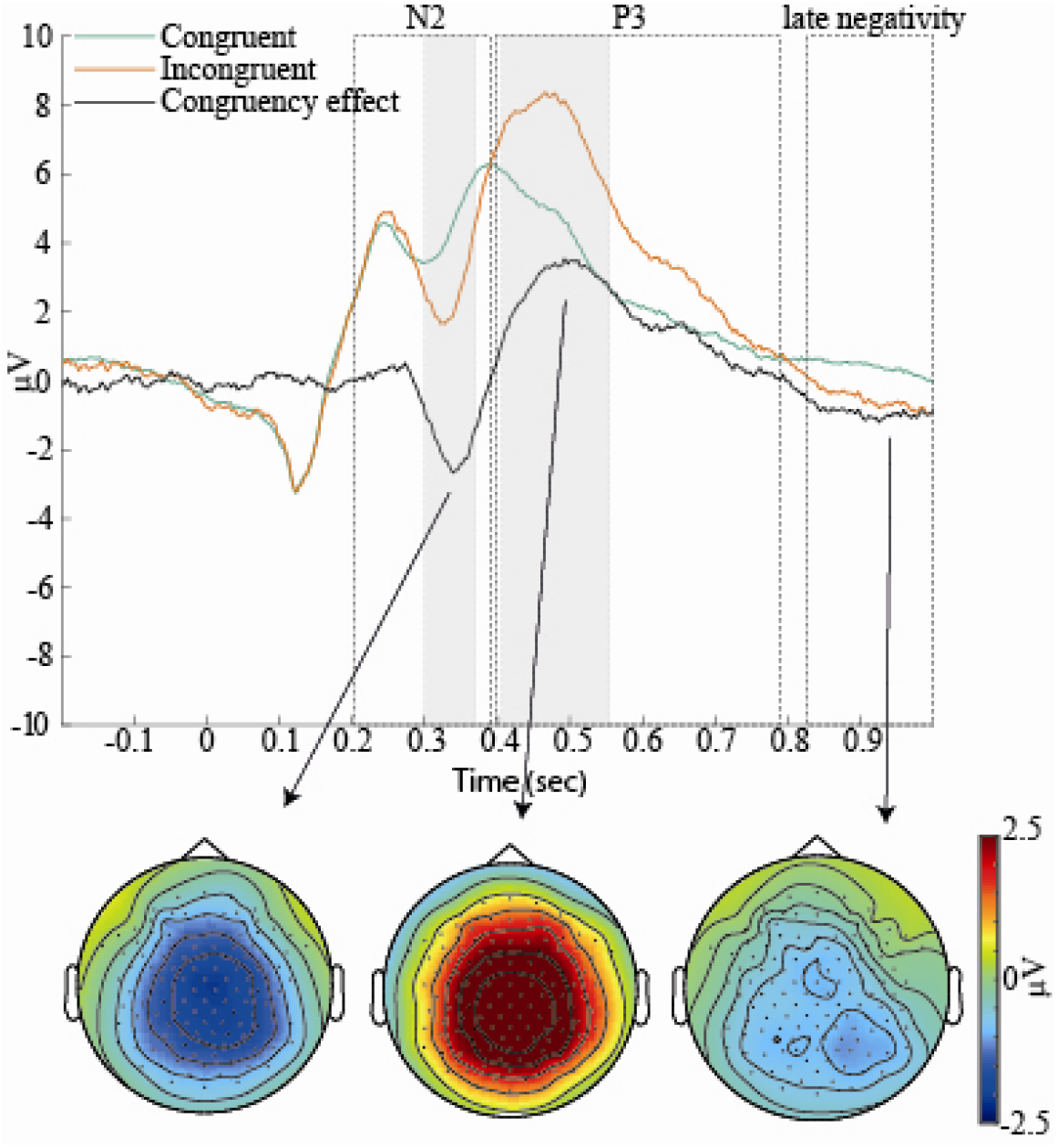
Stimulus locked averaged ERPs for congruent trials (green), incongruent trials (orange) and the congruency effect (black) in the flanker task, averaged across all participants at the FCz channel. The dotted rectangles represent the time windows of the three significant between-condition differences (i.e., the N2, P3, and late negativity). The head plots illustrate the topographic distribution of these differences, averaged over the respective significant time window (note that the grey shading is purely for illustrative purposes centred around the N2 (300-380ms) and P3 (410-550ms) peaks with the topographic distribution matching the respective time windows). The topographic distribution of the late negativity component is averaged over the time window marked by the dotted rectangle.

We observed a significant condition difference (*p = .004*), with the incongruent condition leading to more pronounced negative amplitudes compared to the congruent condition. This corresponded to a cluster ~200 to 400ms post stimulus presentation and was maximal over central sites. This reflected a condition difference at the N2, maximal between ~300 and 380ms. Second, there was also a significant P3 difference (*p < .001*), with the incongruent condition leading to a bigger P3 component compared to the congruent condition. This effect was reflected in a cluster spanning from 390 to 790ms and was maximal over central electrodes. Third, we found a significant condition difference (*p = .004*) with the incongruent condition leading to a greater late negativity (compared to congruent trials). This corresponded to a cluster from 840 to 1000ms, maximal over central (and posterior) sites.

#### ERP language group differences

The results of the non-parametric cluster-based permutation tests of ERP group differences (monolinguals vs. bilinguals) are shown in Figure 3 for each condition (congruent and incongruent conditions) and in Figure 4 for the congruency effect (incongruent condition - congruent condition). We observed significant language group effects in both congruent and incongruent conditions. In the congruent condition, the cluster-based permutation test indicated that there was a significant effect of language group (*p = .018*). We observed a group difference in a time interval encompassing the N2 and P3 components, extending from 220ms - 460ms. Within this time window, the N2 component (defined upon visual inspection between 275-325ms) was maximal over fronto-central sites, and the P3 component (defined upon visual inspection between 375-425ms) was maximal over fronto-central and right centro-parietal electrodes. The N2 was more pronounced (i.e., more pronounced negative-going amplitudes) in bilinguals compared to monolinguals and the P3 was smaller in bilinguals compared to monolinguals. In the incongruent condition, the cluster□based permutation test again indicated that there was a significant effect of language group (*p = .014*). We observed a significant difference in the ERP waveform from around 240ms to 480ms. Within this time window, the N2 component (defined upon visual inspection between 325-375ms) was maximal over fronto-central sites, and the P3 component (defined upon visual inspection between 425-475ms) was maximal over right fronto-temporal sites. Thus in both conditions, the N2 was more pronounced (i.e., greater negative peak) in bilinguals than in monolinguals and the P3 was smaller in bilinguals than in monolinguals. While Figure 4 suggests a smaller congruency effect for P3 amplitudes, the cluster based permutation tests did not reveal any significant group differences.

**Figure 3.**
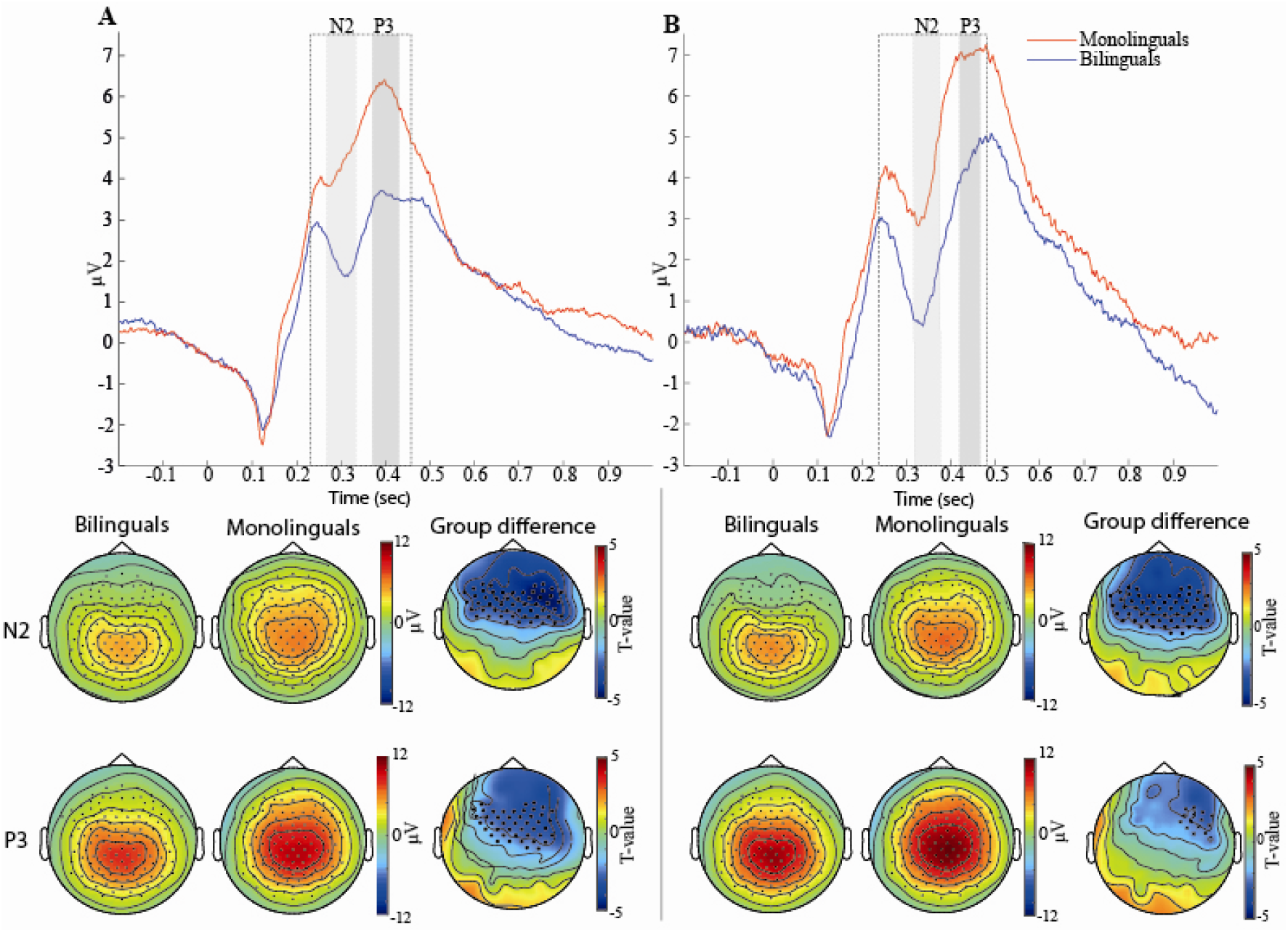
Stimulus locked averaged ERPs produced by (A) congruent trials and (B) incongruent trials in the flanker task for monolinguals (red) and bilinguals (blue). The dotted rectangles represent the time windows of the significant between-group differences for each condition. The ERP waveforms show averaged ERPs across the electrode clusters that indicate the maximal group difference in the specific time interval (this is for illustrative purposes only, the whole time window of 0 to 1 s was used in the non-parametric cluster-based permutation tests). The black dots in the topographic head plots illustrate the significant clusters of electrodes for group differences for the N2 and P3 components. The time windows for N2 and P3 components are shaded in grey and were defined based on visual inspection.

**Figure 4.**
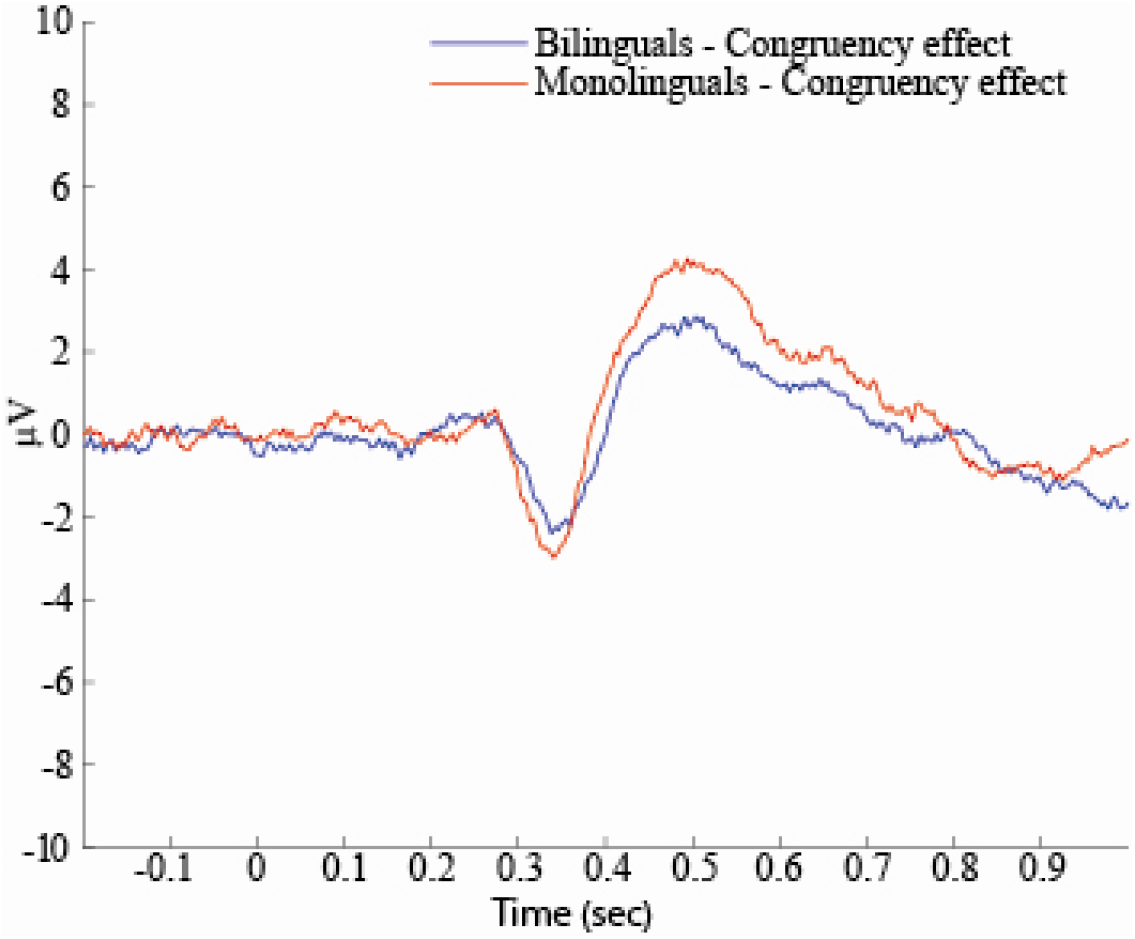
Stimulus locked averaged ERP congruency effect (incongruent trials – congruent trials) for bilinguals (blue) and monolinguals (red) at an exemplary electrode FCz. While monolinguals appear to have a larger congruency effect at the P3 component than bilinguals, this difference was not significant.

#### Correlations between behavioural data and mean peak ERP amplitudes

To investigate whether stronger or weaker engagement of neural processes in the task was beneficial for behavioural performance or not, we correlated the mean peak ERP amplitudes of N2 and P3 for each participant with their ex-Gaussian μ and τ parameters for each condition (congruent and incongruent). Note that mean N2 and P3 peaks here are assumed to be reflections of a participant’s average conflict monitoring and attentional engagement for cognitive control across the task. Ex-Gaussian parameter μ reflects the mean RT of a participant’s majority responses and ex-Gaussian parameter τ their extremeness and the frequency of their slow responses. To see whether any relationship that we found between a neural marker and a behavioural marker differed between groups, we conducted a regression analysis where we checked for an interaction between the ERP component and the language group factor (monolinguals vs. bilinguals). In case of a significant interaction, we conducted follow-up correlation analyses for both groups.

The time windows of the N2 and P3 components were defined upon visual inspection of the ERP data. The time window used for calculating the mean peak amplitude of the N2 component was 275 to 325ms for the congruent condition and 325 to 375 ms for the incongruent condition (see also Figure 3). The time window used for calculating the mean peak amplitude of the P3 component was 375 to 425 ms for the congruent condition and 425 to 475 ms for the incongruent condition (see also Figure 3). The electrodes used for calculating the mean peak amplitudes map onto the electrode clusters that revealed significant group differences in the N2 and P3 for congruent and incongruent conditions (marked as black dots in the topographic plots in Figure 3).

Pearson’s *r* results are shown in Figures 5 and 6 for μ and *τ* respectively. Due to multiple comparisons for each ERP component, we adjusted α to .05/4 = *.0125*. Ex-Gaussian μ did not significantly correlate with any of the two ERP peaks. In contrast, ex-Gaussian *τ* correlated negatively with both N2 and P3 peak amplitudes in both conditions (although the correlation with N2 in the congruent condition was merely a trend with adjusted α). Since some relationships showed outliers, we also calculated Spearman’s ρ. Results were very similar to those for Pearson *r*, apart from a trend for the correlation of *τ* with P3 in the incongruent condition, ρ = −.23, *p =* .098. Follow-up regression analyses did not show significant interactions of language group with neural markers for most correlations (congruent condition: language group x N2, *t*(50) = −.60, *p = .552;* language group x P3, *t*(50) = 1.81, *p = .076*), suggesting that the relationships between the neural markers and τ held for both participant groups. However, the P3 x language group interaction for the incongruent condition was significant (*t*(50) = 3.12, *p = .003*). P3 correlated with τ only for bilinguals (r = −.51, p = .007; Spearman ρ = −.40, p = .040), not for monolinguals (r = −.12, p = .528).

**Figure 5.**
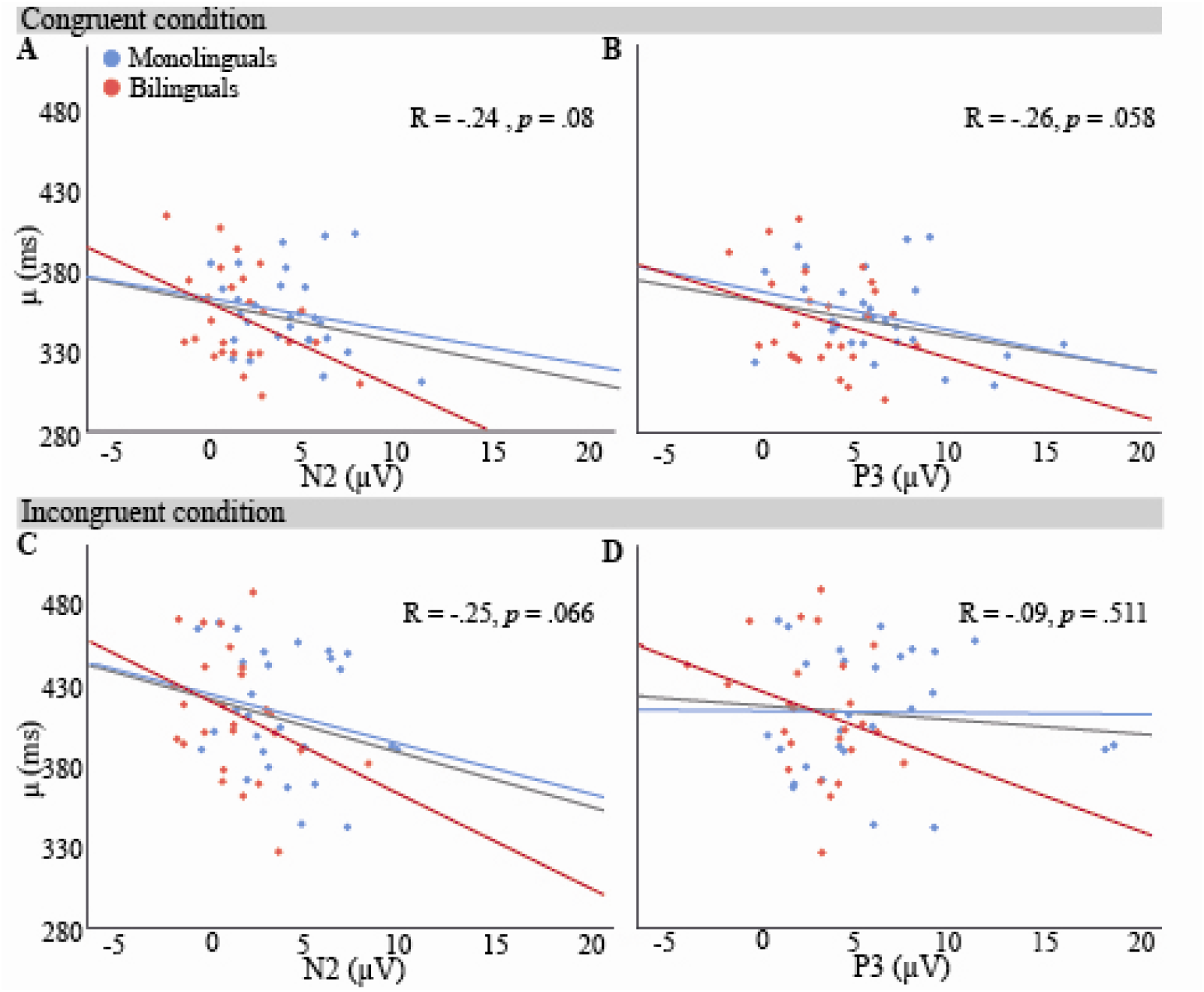
Relationships (Pearson r) of μ (ms) with mean N2 and P3 peak amplitudes (μV) in the congruent condition (panels A and B) and in the incongruent condition (panels C and D), collapsed over participant groups (N=54). None of the relationships were significant (α = .0125). Monolingual data are marked in blue, bilingual data in red. All lines of best fit are least squares regression lines; the grey line reflects all data points, the red line represents bilinguals and the blue line represents monolinguals.

**Figure 6.**
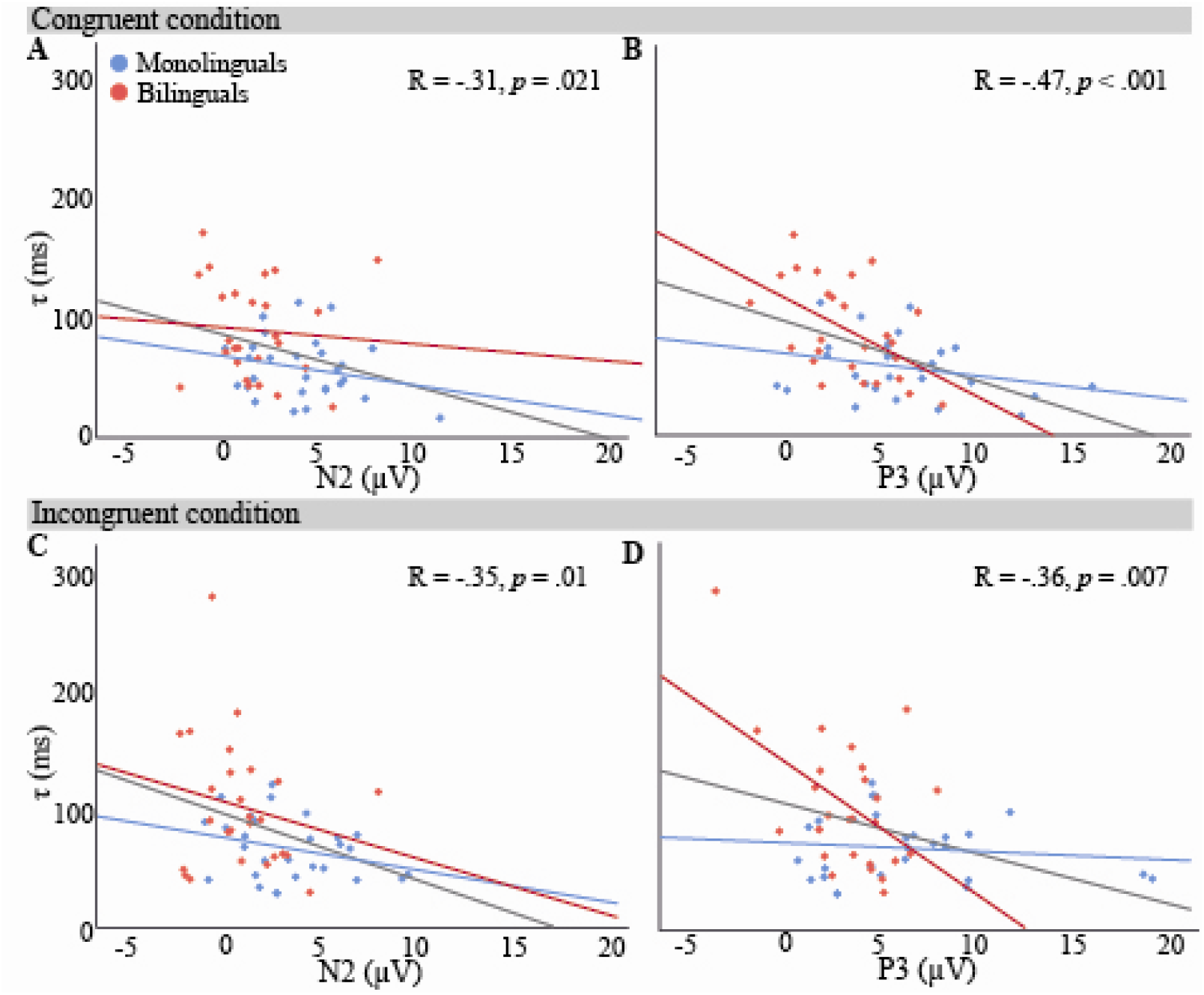
Relationships (Pearson r) of *τ* (ms) with mean N2 and P3 amplitudes (μV) in the congruent condition (panels A and B) and in the incongruent condition (panels C and D), collapsed over participant groups (N = 54). Apart from a trend for a relationship with N2 in the congruent condition, all correlations were significant (α = .0125). Monolingual data are marked in blue, bilingual data in red. All lines of best fit are least squares regression lines; the grey line of best fit reflects all data points, the red line represents bilinguals and the blue line represents monolinguals. Relationships are not affected by language group, apart from the relationship with P3 in the incongruent condition, which was significant only for bilinguals.

## Discussion

In the present study, we aimed to investigate differences in engagement of sub-processes of cognitive control between monolingual and bilingual speakers and how these might lead to behavioural differences. For that we examined the effect of bilingualism on behavioural and neural ERPs responses in a flanker task. We replicated what has been previously observed with regard to behavioural and neural differences for congruent and incongruent conditions. With regards to task performance differences between monolingual and bilingual speakers, we found that bilinguals had longer response distribution tails than monolinguals, independent of flanker condition (reflected also in a trend towards overall slower RTs in bilinguals, see Supplementary material). Bilinguals also exhibited more pronounced N2 and smaller P3 ERP components compared to their monolingual counterparts, independent of condition. We did not find any group differences in terms of congruency effects, neither for behavioural nor for neural responses. Furthermore, behavioural and ERP responses were related in that larger N2 components and smaller P3 components were correlated with longer distribution tails. These relationships were not affected by group, apart from the relationship between P3 and response distribution tails for the incongruent condition, which was present only for bilinguals.

### Behavioural data

As expected, the flanker task produced typical behavioural congruency effects, with incongruent trials yielding longer RTs and lower accuracy rates compared to congruent trials. This reflects the interference from the incongruent flankers that required more conflict monitoring and conflict resolution (Eriksen & Eriksen, 1974). Furthermore, we examined the ex-Gaussian parameters μ and τ of the RT distributions to assess differences in automatic and controlled processes. In line with previous literature (Heathcote et al., 1991; Spieler et al., 1996; Zhou & Krott, 2018), incongruent trials produced larger μ compared to congruent trials (in both language groups), reflecting an anticipated positive shift in response time distribution in the more demanding condition. Similar to the increase in averaged RT, the increased μ corresponds with the additional processing cost associated with resolving conflict and active inhibitory mechanisms in the incongruent condition (Zhou & Krott, 2018). We also found that the incongruent condition was associated with larger *τ* compared to the congruent condition (regardless of language group). The *τ* parameter has been shown to be less affected by condition differences in conflict tasks (Aarts et al., 2009; Heathcote et al., 1991; Spieler et al., 2000), but a condition difference in *τ* was found in, for instance, a flanker task by Zhou and Krott (2018). As discussed, the flanker task is thought to engage various controlled processes: conflict monitoring, the deliberate processing route that involves the stimulus-response translation and control processes needed to merge the outputs of the two processing routes. In this context, *τ* is thought to reflect controlled processes. These processes are more active or might take more time for the incongruent condition. Thus, if they are not as efficient for some incongruent trials, they can lead to occasional slow responses.

In regard to group differences, bilingual and monolingual participants did not differ in terms of the bulk of the responses (reflected by μ), suggesting generally similar efficiency in response conflict resolution across the groups (Zhou & Krott, 2018). However, bilinguals exhibited longer response time distribution tails than monolinguals, independent of condition, meaning they had more particularly long responses and these responses were more extreme than for monolinguals (reflected in τ). In terms of averaged RTs, these longer response distribution tails were reflected in a trend for overall slower responses in bilinguals compared to monolinguals. As *τ* could reflect various controlled processes (conflict monitoring, the deliberate processing route and the merging of the two processing routes), group differences can arise from differences in one or more of these processes. In order to determine what these differences might be, it is useful to take into account group differences in neural patterns and how behavioural measures relate to them.

Note that the finding of longer response distribution tails in bilinguals is contrary to previous findings (Abutalebi et al., 2015; Calabria et al., 2011; Tse & Altarriba, 2012; Zhou & Krott, 2018). It is not clear how these differences occur. But again, differences in experimental paradigms (inclusion of a neutral condition, cues preceding the stimulus arrays, carry-over effects from other tasks, different tasks (Stroop instead of flanker task)) might lead to variations in the engagement of different controlled processes.

### Event related brain potential (ERP) data

In terms of differences between conditions and in line with previous studies (Kousaie & Phillips, 2012, 2017; Purmann et al., 2011; van Veen & Carter, 2002; Wild-Wall et al., 2008), greater conflict monitoring demands emerging from the incongruent trials led to a significantly larger N2 component compared to the congruent condition (regardless of language group) over central sites. The N2 peaked between 320 and 380ms, which is in line with previous results for a very similar version of the flanker task as used here (Yeung et al., 2004). The N2 effect is quite central compared to other studies who reported more fronto-central effects. But the distribution is in line with the finding that the N2 is more central and less frontal when incongruent trials are rather infrequent as in our task (Bartholow et al., 2005). The larger amplitude in the incongruent condition has been associated with enhanced conflict monitoring and attention and the devotion of earlier resources for conflict processing (Bartholow et al., 2005; Folstein & Van Petten, 2008; Grundy, Anderson, et al., 2017; Grützmann et al., 2014; Kousaie & Phillips, 2012; Purmann et al., 2011; Tillman & Wiens, 2011; Yeung & Cohen, 2006), an interpretation supported by the findings that the N2 in the flanker task has been localised to the ACC. Furthermore and as expected, the incongruent condition also elicited greater P3 amplitudes compared to the congruent condition (see, e.g., Kopp et al., 1996; Wild-Wall et al., 2008). Incongruent trials required more attentional resources devoted to the conflict resolution due to the contradictory target and distractor information that led to the activation of competing responses. Thus, categorisation was more effortful in the incongruent condition.

Apart from congruency effects on the N2 and P3 components, we also observed a late negativity for incongruent trials from 840 ms. Jiao et al. (2020) also reported such an effect, but they did not interpret it. Importantly, this effect occurred well after responses were given. It is therefore most likely that it reflects either retrospective monitoring processes or the increase of alertness after a challenging incongruent trial. The hypothesis of retrospective monitoring is in line with its distribution, which mirrors that of the N2.

In regard to language group differences, we found that the N2 component was more pronounced in the bilingual group compared to the monolingual group at frontal-central sites independent of condition (thus for both congruent and incongruent trials). In contrast, the P3 component was significantly reduced for bilinguals compared to monolinguals, again independent of condition. The N2 component in a flanker task has been shown to be sensitive to the monitoring demand (Clayson & Larson, 201; Danielmeier et al., 2009; Folstein & Van Petten, 2008; Grützmann et al., 2014; Purmann et al., 2011; Yeung & Cohen, 2006), suggesting that N2 reflects strategic proactive control (Bugg & Gonthier, 2020). Our pattern of ERP results therefore suggests that bilingual speakers allocated greater resources to conflict monitoring than monolingual speakers, independent of condition. Bilinguals thus utilised a more proactive processing approach than monolinguals (Bartholow et al., 2005; DeLuca et al., 2020; Grundy, Anderson, et al., 2017; Tillman & Wiens, 2011). As a consequence of the proactive approach of bilinguals, fewer attentional resources were devoted for conflict control and categorisation, reflected in the smaller frontal-central P3 component in bilinguals compared to monolinguals. In terms of the processes involved in the flanker task, this means that bilinguals devoted fewer attentional resources to the merging of the two processing routes (direct route and deliberate route).

The group effect on P3 was more frontal than the P3 effect caused by the incongruent flankers and group effects reported in the literature for other types of conflict task (Kousaie, 2012; 2017; Moreno et al., 2014). Its distribution resembled rather a P3a than the standard P3 elicited in the flanker task. Both P3 components have been argued to reflect attentional processes (Polich, 2007). But the P3a occurs about 75-100ms earlier than the standard P3 and, in contrast to the standard P3, is elicited by stimuli that are not attended to (see overview in Polich, 2007). Overall, the P3 group effect in our study therefore resembles more the standard P3. On the other hand, the frontal distribution might indicate a stronger contribution of frontal attentional mechanisms, assumed to underlie also the P3a (Polich, 2007). A smaller P3 component for bilingual speakers might therefore reflect a reduced engagement of frontal attentional processes during stimulus categorisation.

Our ERP results are compatible with various findings for conflict tasks. They are in line with findings for Go/No Go tasks, in which bilinguals had more pronounced N2 amplitudes compared to monolinguals in the No Go condition (despite the lack of behavioural performance differences) (Fernandez et al., 2013, 2014; Moreno et al., 2014), and in which higher L2 proficiency has been found to lead to larger N2 amplitudes (Fernandez et al., 2013, 2014). The combination of increased N2 and reduced P3 in our study is equivalent to what Jiao et al. (2020) had found for flanker trials embedded in a mixed language context, thus in a situation of enhanced executive control demand. Similarly, a reduced P3 in flanker trials had been reported for a mixed compared to a single language condition when participants did not have to produce, but to comprehend the language stimuli (Wu & Thierry, 2013). Furthermore, bilingual speakers have shown reduced P3 amplitudes compared to monolingual speakers in a Flanker task (Botezatu et al., 2021) previously, as well as in a Stroop task (Coderre et al., 2014) and a Simon task (Kousaie & Phillips, 2012). These results all suggest that bilinguals allocate less resources to the merging of competing responses and categorisation processes than monolingual speakers in a conflict task.

In contrast, Kousaie and Phillips (2012, 2017) found no or opposite patterns of group differences in ERPs. They did not find language group differences in N2 or P3 amplitudes in a flanker task. Instead, they discovered the opposite N2 effect in a verbal Stroop task, with monolinguals showing a larger N2 than bilinguals. As noted, the results for the verbal Stroop task might be due to more resources being diverted to linguistic processes during the task. Finally, bilinguals showed greater instead of reduced P3 amplitudes in Stroop and Simon tasks. One remarkable difference between these two studies and the current study is that they included not only congruent and incongruent trials, but also 33% neutral trials. Since both N2 and P3 are very sensitive to executive function demands of a task, especially monitoring demands (Jiao et al., 2020; Purmann et al., 2011; Wu & Thierry, 2013), it is possible that the inclusion of neutral trials shifted the balance of monitoring and attention to conflict resolution processes in the two participant groups. While the proportion of incongruent trials (25% in the current version and 33% in Kousaie and Philips’ version) is quite similar, participants had to only monitor for incongruent flankers in Kousaie and Philips’ studies when a stimulus actually included flankers (i.e. for incongruent and congruent trials). This might have changed bilinguals’ proactive monitoring approach to a more reactive approach, thus one very similar to that of monolingual participants. This also explains that Botezatu et al. (2021) did not find an N2 group difference when using the same proportion of trials as Kousaie and Philips.

Proactive control is more effortful, but known to be generally more effective than reactive control (Braver, 2012) and therefore usually advantageous. However, bilinguals’ proactive enhanced resource allocation to monitoring was not quite advantageous in the present study. As larger N2 amplitudes were related with longer distribution tails, their enhanced monitoring occasionally led to very slow responses. Stronger engagement of monitoring processes might mean that responses were checked particularly thoroughly, for instance by increasing a response activation threshold. This would mean that bilinguals over-engaged in monitoring in the current study. As mentioned, the relatively rare (25%) occurrence of incongruent trials might have activated the generally proactive monitoring approach in bilinguals, while monolinguals took a more reactive processing approach. The latter might have been slightly more beneficial, as evidenced in fewer extremely long responses in the monolingual group. In other words, bilinguals’ more efficient conflict resolution and categorisation due to reduced attentional resources devoted to the merging of the two processing routes (reflected in reduced P3 amplitudes) was hampered by a hyper-activate monitoring system in the present task.

Relationships between neural measures (N2 and P3) and behavioural measures (ex-Gaussian parameters) were generally not affected by participant group membership, meaning that processes associated with the N2 and P3 seem to be very similar for both groups. However, the P3 was related to response distribution tails in the incongruent condition for bilinguals only. Thus, the up- and down-regulating of attentional resources for conflict processing seems to be linked to the occurrence of long responses in bilinguals only. It is not clear why this might be the case. As correlations can be affected by sample size, future studies will need to investigate whether this can be replicated with a larger participant group.

### The bilingual advantage in conflict tasks

What do the current findings mean with regard to the debate about the bilingual advantage in conflict tasks? As mentioned, the current bilingual literature is equivocal in regard to language group differences in such tasks. Specifically, in the flanker task, some previous studies have found that bilinguals are faster (Costa et al., 2008; Emmorey et al., 2008), and more accurate (Kousaie & Phillips, 2017) than monolinguals. Other studies have reported no significant RT differences between bilinguals and monolinguals (Antón et al., 2019; Grundy, Chung-Fat-Yim, et al., 2017; Kousaie & Phillips, 2012, 2017; Luk et al., 2010) or (similarly to our study) a small non-significant bilingual disadvantage (Paap & Greenberg, 2013; Paap & Sawi, 2014). Together these results suggest that there is no consistent advantage in flanker tasks, or more generally in conflict tasks (see Ware et al., 2020, who found that different tasks are more or less consistent in showing a bilingual advantage). An advantage likely only appears in conditions with the right balance of demands on sub-processes. If proactive monitoring is advantageous in a paradigm, bilinguals might process the conflict more efficiently and therefore more likely show a behavioural advantage. But, proactive conflict monitoring does not always lead to faster (or more accurate) responses as we have seen in the current study, it can also backfire and hinder performance.

### Individual differences

While group differences in conflict tasks can be affected by task parameters, performance of bilinguals is also impacted by individual variations in bilingual experience. Language experiences such as intensity and diversity of L1 and L2 use, language switching in daily life, relative proficiency and duration of the two languages have varying consequences on control demands and lead to both structural and functional neural changes (see the Unifying the Bilingual Experience Trajectories (UBET) model by DeLuca et al., 2020). For instance, models of individual variation of bilingual experience onto structural and functional brain changes predict less reliance on frontal areas with longer L2 use (DeLuca et al., 2020; Grundy, Anderson, et al., 2017). Not surprisingly, there are increasing calls to account for individual differences in bilingual experiences in the bilingual advantage literature (Poarch & Krott, 2019). The effect of such variations has indeed been shown in both the flanker task (Dong & Zhong, 2017; Hofweber et al., 2016) and in other tasks with a suppression component (Fernandez et al., 2013, 2014; Gullifer et al., 2018; Sullivan et al., 2014). For the flanker task, bilinguals who engage in more dense code-switching showed inhibitory advantages in a condition with medium monitoring demand (25% incongruent trials) (Hofweber et al., 2016); and interpreting experience heightens early attentional processing (larger N1), conflict monitoring (larger N2), and interference suppression (smaller P3 and smaller RT interference effect) (Dong & Zhong, 2017). L2 proficiency seems to particularly impact neural markers in a Go/No Go task, with higher proficiency leading to more pronounced N2 (Fernandez et al., 2013) and a six-month University Spanish course (compared to a Psychology course) increasing the P3 component (Sullivan et al., 2014). Furthermore, proactive and reactive control can be affected by different experiences. Gullifer et al. (2018) reported that greater diversity in language use in daily life was related with greater reliance on proactive control in an AX-Continuous performance task, while earlier L2 age of acquisition was associated with a decrease in reliance on proactive control. Future studies will therefore need to take into account not only task parameters, but also individual variations in bilingual experience.

To conclude, we have found that bilingual speakers showed evidence for enhanced proactive monitoring during a flanker task with medium monitoring demand compared to monolingual speakers. The enhanced monitoring was followed by less attentional resources devoted to conflict resolution and stimulus categorisation, thus to less effortful categorisation. The monitoring system, however, was rather overactive as it led occasionally to very slow responses. Thus, while the processes of monitoring and categorisation more or less balanced each other out, the overactive and therefore less efficient monitoring slightly dominated. These results demonstrate how the engagement of sub-processes of a task together determine overall behavioural performance and can affect group differences. While there is currently a strong focus shift in the literature on how individual differences in bilingual experience might affect bilingual performance and therefore differences compared to monolingual speakers, we propose that the study of balance of sub-processes in conflict tasks is a complementary avenue that is useful for a better understanding of any functional differences between bilingual and monolingual speakers.

## References

Aarts, E., Roelofs, A., & van Turennout, M. (2009). Attentional control of task and response in lateral and medial frontal cortex: Brain activity and reaction time distributions. Neuropsychologia, 47(10), 2089–2099. https://doi.org/10.1016/j.neuropsychologia.2009.03.019

Abutalebi, J., Guidi, L., Borsa, V., Canini, M., Della Rosa, P. A., Parris, B. A., & Weekes, B. S. (2015). Bilingualism provides a neural reserve for aging populations. Neuropsychologia, 69, 201–210. https://doi.org/10.1016/j.neuropsychologia.2015.01.040

Antón, E., Carreiras, M., & Duñabeitia, J. A. (2019). The impact of bilingualism on executive functions and working memory in young adults. PLoS ONE, 14(2), 1–30. https://doi.org/10.1371/journal.pone.0206770

Barac, R., Moreno, S., & Bialystok, E. (2016). Behavioral and Electrophysiological Differences in Executive Control Between Monolingual and Bilingual Children. Child Development, 87(4), 1277–1290. https://doi.org/10.1111/cdev.12538

Bartholow, B. D., Pearson, M. A., Dickter, C. L., Sher, K. J., Fabiani, M., & Gratton, G. (2005). Strategic control and medial frontal negativity □: Beyond errors and response conflict. Psychophysiology, 42, 33–42. https://doi.org/10.1111/j.1469-8986.2005.00258.x

Bialystok, E., Craik, F. I. M., & Luk, G. (2012). Bilingualism: consequences for mind and brain. Trends in Cognitive Sciences, 16(4), 240–250. https://doi.org/10.1016/j.tics.2012.03.001

Bialystok, E., Klein, R., Craik, F. I. M., & Viswanathan, M. (2004). Bilingualism, aging, and cognitive control: Evidence from the Simon task. Psychology and Aging, 19(2), 290–303. https://doi.org/10.1037/0882-7974.19.2.290

Botezatu, M. R., Miller, C. A., Johnson, J., & Misra, M. (2021). Event-related potentials reveal that bilinguals are more efficient in resolving conflict than monolinguals. NeuroReport, 32(8), 721–726. https://doi.org/10.1097/WNR.0000000000001645

Botvinick, M. M., Cohen, J. D., & Carter, C. S. (2004). Conflict monitoring and anterior cingulate cortex: An update. Trends in Cognitive Sciences, 8(12), 539–546. https://doi.org/10.1016/j.tics.2004.10.003

Braver, T. S. (2012). The variable nature of cognitive control: A dual-mechanisms framework Shifting the emphasis to variability in cognitive control. Trends in Cognitive Sciences, 16(2), 106–113. https://doi.org/10.1016/j.tics.2011.12.010.The

Brown, S., & Heathcote, A. (2003). QMLE: Fast, robust, and efficient estimation of distribution functions based on quantiles. Behavior Research Methods, Instruments, and Computers, 35(4), 485–492. https://doi.org/10.3758/BF03195527

Calabria, M., Hernández, M., Martin, C. D., & Costa, A. (2011). When the tail counts: the advantage of bilingualism through the ex-gaussian distribution analysis. Frontiers in Psychology, 2(250), 1–8. https://doi.org/10.3389/fpsyg.2011.00250

Carter, C. S., Braver, T. S., Barch, D. M., Botvinick, M. M., Noll, D., & Cohen, J. D. (1998). Anterior cingulate cortex, error detection, and the online monitoring of performance. Science, 280(5364), 747–749. https://doi.org/10.1126/science.280.5364.747

Carter, C. S., & van Veen, V. (2007). Anterior cingulate cortex and conflict detection: An update of theory and data. Cognitive, Affective, & Behavioral Neuroscience, 7(4), 367–379. https://doi.org/10.3758/CABN.7.4.367

Chung-Fat-Yim, A., Poarch, G. J., Comishen, K. J., & Bialystok, E. (2021). Does language context impact the neural correlates of executive control in monolingual and multilingual young adults? Brain and Language, 222(105011), 1–12. https://doi.org/10.1016/j.bandl.2021.105011

Clayson, P. E., & Larson, M. J. (2011). Conflict adaptation and sequential trial effects: Support for the conflict monitoring theory. Neuropsychologia, 49(7), 1953–1961. https://doi.org/https://doi.org/10.1016/j.neuropsychologia.2011.03.023

Coderre, E. L., Walter, J. B., & Heuven, V. (2014). Electrophysiological explorations of the bilingual advantage: Evidence from a Stroop task. PLoS ONE, 9(7). https://doi.org/10.1371/journal.pone.0103424

Coles, M. G. H., Gratton, G., Bashore, T. R., Eriksen, C. W., & Donchin, E. (1985). A Psychophysiological Investigation of the Continuous Flow Model of Human Information Processing. Journal of Experimental Psychology: Human Perception and Performance, 11(5), 529–553. https://doi.org/10.1037/0096-1523.11.5.529

Costa, A., Hernández, M., Costa-Faidella, J., & Sebastián-Gallés, N. (2009). On the bilingual advantage in conflict processing: Now you see it, now you don’t. Cognition, 113(2), 135–149. https://doi.org/10.1016/j.cognition.2009.08.001

Costa, A., Hernández, M., & Sebastián-Gallés, N. (2008). Bilingualism aids conflict resolution: Evidence from the ANT task. Cognition, 106(1), 59–86. https://doi.org/10.1016/j.cognition.2006.12.013

Danielmeier, C., Wessel, J. R., Steinhauser, M., & Ullsperger, M. (2009). Modulation of the error-related negativity by response conflict. Psychophysiology, 46(6), 1288–1298. https://doi.org/10.1111/j.1469-8986.2009.00860.x

Delorme, A., & Makeig, S. (2004). EEGLAB: An open source toolbox for analysis of single-trial EEG dynamics including independent component analysis. Journal of Neuroscience Methods, 134(1), 9–21. https://doi.org/10.1016/j.jneumeth.2003.10.009

DeLuca, V., Segaert, K., Mazaheri, A., & Krott, A. (2020). Understanding bilingual brain function and structure changes? U Bet! A Unified Bilingual Experience Trajectory model. Journal of Neurolinguistics, 56(2020), 1–14. https://doi.org/10.1017/CBO9781107415324.004

Dong, Y., & Zhong, F. (2017). Interpreting experience enhances early attentional processing, conflict monitoring and interference suppression along the time course of processing. Neuropsychologia, 95, 193–203. https://doi.org/10.1016/j.neuropsychologia.2016.12.007

Donnelly, S., Brooks, P. J., & Homer, B. D. (2019). Is there a bilingual advantage on interference-control tasks? A multiverse meta-analysis of global reaction time and interference cost. Psychonomic Bulletin and Review, 26(4), 1122–1147. https://doi.org/10.3758/s13423-019-01567-z

Duñabeitia, J. A., Hernández, J. A., Antón, E., Macizo, P., Estévez, A., Fuentes, L. J., & Carreiras, M. (2014). The inhibitory advantage in bilingual children revisited: Myth or reality? Experimental Psychology, 61(3), 234–251. https://doi.org/10.1027/1618-3169/a000243

Emmorey, K., Luk, G., Pyers, J. E., & Bialystok, E. (2008). The source of enhanced cognitive control in bilinguals: Evidence from bimodal bilinguals. Psychological Science, 19(12), 1201–1206. https://doi.org/10.1111/j.1467-9280.2008.02224.x

Epstein, J. N., Langberg, J. M., Rosen, P. J., Graham, A., Narad, M. E., Antonini, T. N., Brinkman, W. B., Froehlich, T., Simon, J. O., & Altaye, M. (2011). Evidence for Higher Reaction Time Variability for Children With ADHD on a Range of Cognitive Tasks Including Reward and Event Rate Manipulations. Neuropsychology, 25(4), 427–441. https://doi.org/10.1037/a0022155

Eriksen, B. A., & Eriksen, C. W. (1974). Effects of noise letters upon the identification of a target letter in a nonsearch task. Perception & Psychophysics, 16(1), 143–149. https://doi.org/10.3989/arbor.2000.i650.965

Eriksen, C. W., & Schultz, D. W. (1979). Information processing in visual search: A continuous flow conception and experimental results. Perception & Psychophysics, 25(4), 249–263. https://doi.org/10.3758/BF03198804

Fernandez, M., Acosta, J., Douglass, K., Doshi, N., & Tartar, J. L. (2014). Speaking two languages enhances an auditory but not a visual neural marker of cognitive inhibition. AIMS Neuroscience, 1(2), 145–157. https://doi.org/10.3934/Neuroscience.2014.2.145

Fernandez, M., Tartar, J. L., Padron, D., & Acosta, J. (2013). Neurophysiological marker of inhibition distinguishes language groups on a non-linguistic executive function test. Brain and Cognition, 83(3), 330–336. https://doi.org/10.1016/j.bandc.2013.09.010

Folstein, J. R., & Van Petten, C. (2008). Influence of cognitive control and mismatch on the N2 component of the ERP: A review. Psychophysiology, 45(1), 152–170. https://doi.org/10.1111/j.1469-8986.2007.00602.x

Forster, S. E., Carter, C. S., Cohen, J. D., & Cho, R. Y. (2011). Parametric manipulation of the conflict signal and control-state adaptation. Journal of Cognitive Neuroscience, 23(4), 923–935. https://doi.org/10.1162/jocn.2010.21458

Gathercole, V. C. M., Thomas, E. M., Kennedy, I., Prys, C., Young, N., Guasch, N. V., Roberts, E. J., Hughes, E. K., & Jones, L. (2014). Does language dominance affect cognitive performance in bilinguals? Lifespan evidence from preschoolers through older adults on card sorting, Simon, and metalinguistic tasks. Frontiers in Psychology, 5(11), 1–14. https://doi.org/10.3389/fpsyg.2014.00011

Gratton, G., Coles, M. G. H., & Donchin, E. (1992). Optimizing the use of information: Strategic control of activation of responses. Journal of Experimental Psychology., 121(4), 480–506.

Gratton, G., Coles, M. G., Sirevaag, E. J., Eriksen, C. W., & et al. (1988). Pre- and poststimulus activation of response channels: A psychophysiological analysis. Journal of Experimental Psychology: Human Perception and Performance, 14(3), 331–344. https://doi.org/10.1037//0096-1523.14.3.331

Grundy, J. G. (2020). The effects of bilingualism on executive functions: an updated quantitative analysis. Journal of Cultural Cognitive Science, 4(2), 177–199. https://doi.org/10.1007/s41809-020-00062-5

Grundy, J. G., Anderson, J. A. E., & Bialystok, E. (2017). Neural correlates of cognitive processing in monolinguals and bilinguals. Annals of the New York Academy of Sciences, February. https://doi.org/10.1111/nyas.13333

Grundy, J. G., Chung-Fat-Yim, A., Friesen, D. C., Mak, L., & Bialystok, E. (2017). Sequential congruency effects reveal differences in disengagement of attention for monolingual and bilingual young adults. Cognition, 163, 42–55. https://doi.org/10.1016/j.cognition.2017.02.010

Grützmann, R., Riesel, A., Klawohn, J., Kathmann, N., & Endrass, T. (2014). Complementary modulation of N2 and CRN by conflict frequency. Psychophysiology, 51, 761–772. https://doi.org/10.1111/psyp.12222

Gullifer, J. W., Chai, X. J., Whitford, V., Pivneva, I., Baum, S., Klein, D., & Titone, D. (2018). Bilingual experience and resting-state brain connectivity: Impacts of L2 age of acquisition and social diversity of language use on control networks. Neuropsychologia, 117(2018), 123–134. https://doi.org/10.1016/j.neuropsychologia.2018.04.037

Heathcote, A., Popiel, S. J., & Mewhort, D. J. K. (1991). Analysis of response time distributions: an example using the stroop task. Psychological Bulletin, 109(2), 340–347. https://doi.org/10.1037/0033-2909.109.2.340

Hernández, M., Costa, A., Fuentes, L. J., Vivas, A. B., & Sebastián-Gallés, N. (2010). The impact of bilingualism on the executive control and orienting networks of attention. Bilingualism: Language and Cognition, 13(3), 315–325. https://doi.org/10.1017/S1366728909990010

Hervey, A. S., Epstein, J. N., Curry, J. F., Tonev, S., Eugene Arnold, L., Keith Conners, C., Hinshaw, S. P., Swanson, J. M., & Hechtman, L. (2006). Reaction time distribution analysis of neuropsychological performance in an ADHD sample. Child Neuropsychology, 12(2), 125–140. https://doi.org/10.1080/09297040500499081

Hofweber, J., Marinis, T., & Treffers-Daller, J. (2016). Effects of dense code-switching on executive control. Linguistic Approaches to Bilingualism, 6(5), 648–668. https://doi.org/10.1075/sibil.57.11hof

Jiao, L., Grundy, J. G., Liu, C., & Chen, B. (2020). Language context modulates executive control in bilinguals: Evidence from language production. Neuropsychologia, 142(19). https://doi.org/10.1016/j.neuropsychologia.2020.107441

Johnson, R. J. (1993). On the neural generators of the P300 component of the event-related potential. Psychophysiology, 30, 90–97.

Kałamała, P., Walther, J., Zhang, H., Diaz, M., Senderecka, M., & Wodniecka, Z. (2022). The use of a second language enhances the neural efficiency of inhibitory control: An ERP study. Bilingualism: Language and Cognition, 25(1), 163–180. https://doi.org/10.1017/S1366728921000389

Kane, M. J., & Engle, R. W. (2003). Working-Memory Capacity and the Control of Attention: The Contributions of Goal Neglect, Response Competition, and Task Set to Stroop Interference. Journal of Experimental Psychology: General, 132(1), 47–70. https://doi.org/10.1037/0096-3445.132.1.47

Kok, A. (2001). On the utility of P3 amplitude as a measure of processing capacity. Psychophysiology, 38(3), 557–577. https://doi.org/10.1017/S0048577201990559

Kopp, B., Fred, R., & Mattler, U. (1996). N200 in the flanker task as a neurobiological tool for investigating executive control. Psychophysiology, 33, 282–294.

Kousaie, S., & Phillips, N. A. (2012). Conflict monitoring and resolution: Are two languages better than one? Evidence from reaction time and event-related brain potentials. Brain Research, 1446(2012), 71–90. https://doi.org/10.1016/j.brainres.2012.01.052

Kousaie, S., & Phillips, N. A. (2017). A behavioural and electrophysiological investigation of the effect of bilingualism on aging and cognitive control. Neuropsychologia, 94(July 2016), 23–35. https://doi.org/10.1016/j.neuropsychologia.2016.11.013

Lehtonen, M., Soveri, A., Laine, A., Järvenpää, J., de Bruin, A., & Antfolk, J. (2018). Is bilingualism associated with enhanced executive functioning in adults? A meta-analytic review. Psychological Bulletin, 144(4), 394–425. https://doi.org/10.1037/bul0000142

Li, P., Legault, J., & Litcofsky, K. A. (2014). Neuroplasticity as a function of second language learning: Anatomical changes in the human brain. Cortex, 58, 301–324. https://doi.org/10.1016/j.cortex.2014.05.001

Luk, G., Anderson, J. A. E., Craik, F. I. M., Grady, C. L., & Bialystok, E. (2010). Distinct neural correlates for two types of inhibition in bilinguals: Response inhibition versus interference suppression. Brain and Cognition, 74(3), 347–357. https://doi.org/10.1016/j.bandc.2010.09.004

Maris, E., & Oostenveld, R. (2007). Nonparametric statistical testing of EEG- and MEG-data. Journal of Neuroscience Methods, 164(1), 177–190. https://doi.org/10.1016/j.jneumeth.2007.03.024

Matzke, D., & Wagenmakers, E. J. (2009). Psychological interpretation of the ex-gaussian and shifted wald parameters: A diffusion model analysis. Psychonomic Bulletin and Review, 16(5), 798–817. https://doi.org/10.3758/PBR.16.5.798

Morales, J., Yudes, C., Gomez-Ariza, C. J., & Bajo, M. T. (2015). Bilingualism modulates dual mechanisms of cognitive control: Evidence from ERPs. Neuropsychologia, 66, 157–169. https://doi.org/10.1016/j.neuropsychologia.2014.11.014

Moreno, S., Wodniecka, Z., Tays, W., Alain, C., & Bialystok, E. (2014). Inhibitory Control in Bilinguals and Musicians: Event Related Potential (ERP) Evidence for Experience-Specific Effects. PLoS ONE, 9(4), e94169. https://doi.org/10.1371/journal.pone.0094169

Naeem, K., Filippi, R., Periche-Tomas, E., Papageorgiou, A., & Bright, P. (2018). The importance of socioeconomic status as a modulator of the bilingual advantage in cognitive ability. Frontiers in Psychology, 9(1818), 1–9. https://doi.org/10.3389/fpsyg.2018.01818

Nichols, E. S., Wild, C. J., Stojanoski, B., Battista, M. E., & Owen, A. M. (2020). Bilingualism Affords No General Cognitive Advantages: A Population Study of Executive Function in 11,000 People. Psychological Science, 31(5), 548–567. https://doi.org/10.1177/0956797620903113

Oldfield, R. C. (1971). The assessment and analysis of handedness: The Edinburgh inventory. Neuropsychologia, 9, 97–113. https://doi.org/10.1007/978-0-387-79948-3_6053

Oostenveld, R., Fries, P., Maris, E., & Schoffelen, J. M. (2011). FieldTrip: Open source software for advanced analysis of MEG, EEG, and invasive electrophysiological data. Computational Intelligence and Neuroscience, 2011, 1–9. https://doi.org/10.1155/2011/156869

Oostenveld, R., & Praamstra, P. (2001). The five percent electrode system for high-resolution EEG and ERP measurements. Clinical Neurophysiology, 112(4), 713–719. https://doi.org/10.1016/S1388-2457(00)00527-7

Paap, K. R., & Greenberg, Z. I. (2013). There is no coherent evidence for a bilingual advantage in executive processing. Cognitive Psychology, 66(2), 232–258. https://doi.org/10.1016/j.cogpsych.2012.12.002

Paap, K. R., Johnson, H. A., & Sawi, O. (2015). Bilingual advantages in executive functioning either do not exist or are restricted to very specific and undetermined circumstances. Cortex, 69, 265–278. https://doi.org/10.1016/j.cortex.2015.04.014

Paap, K. R., & Sawi, O. (2014). Bilingual advantages in executive functioning: problems in convergent validity, discriminant validity, and the identification of the theoretical constructs. Frontiers in Psychology, 5(962), 1–15. https://doi.org/10.3389/fpsyg.2014.00962

Pliatsikas, C., & Luk, G. (2016). Executive control in bilinguals: A concise review on fMRI studies. Bilingualism: Language and Cognition, 19(4), 699–705. https://doi.org/10.1017/S1366728916000249

Poarch, G. J., & Krott, A. (2019). A bilingual advantage? An appeal for a change in perspective and recommendations for future research. Behavioral Sciences, 9(9), 1–9. https://doi.org/10.3390/bs9090095

Polich, J. (2007). Updating P300: An integrative theory of P3a and P3b. Clinical Neurophysiology, 118(10), 2128–2148. https://doi.org/10.1016/j.clinph.2007.04.019

Purmann, S., Badde, S., Luna-Rodriguez, A., & Wendt, M. (2011). Adaptation to frequent conflict in the eriksen flanker task: An ERP study. Journal of Psychophysiology, 25, 50–59.

Raven, J. C. (1958). Standard Progressive Matrices. Sets A, B, C, D, and E. London: H.K. Lewis & Co. Ltd.

Ridderinkhof, K. R., van der Molen, M. W., & Bashore, T. R. (1995). Limits on the application of additive factors logic: Violations of stage robustness suggest a dual-process architecture to explain flanker effects on target processing. Acta Psycholoica, 90(1995), 29–48. https://doi.org/https://doi.org/10.1016/0001-6918(95)00031-o

Ridderinkhof, K. R., Wylie, S. A., van den Wildenberg, W. P. M., Bashore, T. R., & van der Molen, M. W. (2021). The arrow of time: Advancing insights into action control from the arrow version of the Eriksen flanker task. Attention, Perception, and Psychophysics, 83(2), 700–721. https://doi.org/10.3758/s13414-020-02167-z

Spieler, D. H., Balota, D. A., & Faust, M. E. (1996). Stroop Performance in Healthy Younger and Older Adults and in Individuals with Dementia of the Alzheimer’s Type. Journal of Experimental Psychology: Human Perception and Performance, 22(2), 461–479. https://doi.org/10.1037/0096-1523.22.2.461

Spieler, D. H., Balota, D. A., & Faust, M. E. (2000). Levels of selective attention revealed through analyses of response time distributions. Journal of Experimental Psychology: Human Perception and Performance, 26(2), 506–526. https://doi.org/10.1037/0096-1523.26.2.506

Sullivan, M. D., Janus, M., Moreno, S., Astheimer, L., & Bialystok, E. (2014). Early stage second-language learning improves executive control: Evidence from ERP. Brain and Language, 139, 84–98. https://doi.org/10.1016/j.bandl.2014.10.004

Tillman, C. M., & Wiens, S. (2011). Behavioral and ERP indices of response conflict in Stroop and flanker tasks. Psychophysiology, 48, 1405–1411. https://doi.org/10.1111/j.1469-8986.2011.01203.x

Tse, C. S., & Altarriba, J. (2012). The effects of first- and second-language proficiency on conflict resolution and goal maintenance in bilinguals: Evidence from reaction time distributional analyses in a Stroop task. Bilingualism: Language and Cognition, 15(3), 663–676. https://doi.org/10.1017/S1366728912000077

Veen, V. Van, & Carter, C. S. (2002). The Timing of Action-Monitoring Processes in the Anterior Cingulate Cortex. Journal of Cognitive Neuroscience, 14(4), 593–602.

Ware, A. T., Kirkovski, M., & Lum, J. A. G. (2020). Meta-Analysis Reveals a Bilingual Advantage That Is Dependent on Task and Age. Frontiers in Psychology, 11(1458), 1–21. https://doi.org/10.3389/fpsyg.2020.01458

Wickens, C., Kramer, A., Vanasse, L., & Donchin, E. (1983). Performance of concurrent tasks: A psychophysiological analysis of the reciprocity of information-processing resources. Science, 221(4615), 1080–1082. https://doi.org/10.1126/science.6879207

Wild-Wall, N., Falkenstein, M., & Hohnsbein, J. (2008). Flanker interference in young and older participants as reflected in event-related potentials. Brain Research, 1211, 72–84. https://doi.org/10.1016/j.brainres.2008.03.025

Wu, Y. J., & Thierry, G. (2013). Fast modulation of executive function by language context in bilinguals. Journal of Neuroscience, 33(33), 13533–13537. https://doi.org/10.1523/JNEUROSCI.4760-12.2013

Yeung, N., Botvinick, M. M., & Cohen, J. D. (2004). The neural basis of error detection: Conflict monitoring and the error-related negativity. Psychological Review, 111(4), 931–959. https://doi.org/10.1037/0033-295X.111.4.931

Yeung, N., & Cohen, J. D. (2006). The Impact of Cognitive Deficits on Conflict Monitoring. Psychological Science, 17(2), 164–171. http://search.ebscohost.com/login.aspx?direct=true&db=aph&AN=19473526&site=ehost-live

Yeung, N., & Nieuwenhuis, S. (2009). Dissociating response conflict and error likelihood in anterior cingulate cortex. Journal of Neuroscience, 29(46), 14506–14510. https://doi.org/10.1523/JNEUROSCI.3615-09.2009

Zhou, B., & Krott, A. (2016). Data trimming procedure can eliminate bilingual cognitive advantage. Psychonomic Bulletin & Review, 23(4), 1221–1230. https://doi.org/10.3758/s13423-015-0981-6

Zhou, B., & Krott, A. (2018). Bilingualism enhances attentional control in non-verbal conflict tasks – evidence from ex-Gaussian analyses. Bilingualism: Language and Cognition, 21(1), 162–180. https://doi.org/10.1017/S1366728916000869

