## Supplementary material for "Bilingual speakers’ enhanced monitoring can slow them down"

**Results of traditional behavioural measures (i.e., averaged RTs and accuracy) on the flanker task for bilingual and monolingual participants**

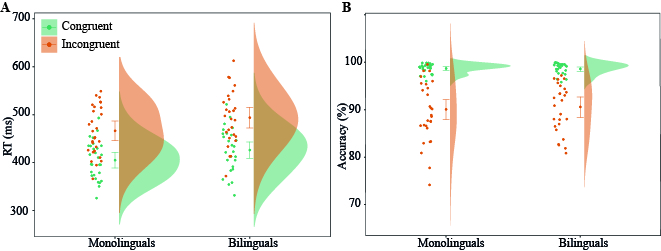

*Supplementary Figure 1.* Distributions and means of RT (panel A), and accuracy (panel B) per condition (congruent and incongruent) in the flanker task for both monolinguals and bilinguals. Error bars represent 95% confidence intervals.

***RT.*** We found a significant main effect of Condition, *F*(1,52) = 405.32, *p < .001*, indicating overall slower RTs in the incongruent compared to the congruent condition. There was a trend for a main effect of Language Group, *F*(1,52) = 3.53, *p = .066*, with bilinguals tending to have overall longer RTs compared to monolinguals. The interaction between Condition and Language Group was not significant, *F*(1,52) = .98, *p = .327*.

***Accuracy.*** There was a significant main effect of Condition, *F*(1,52) = 142.63, *p < .001,* indicating overall lower accuracy in the incongruent compared to the congruent condition. There was no main effect of Language Group, *F*(1,52) = .05, *p =.831,* nor a Language Group by Condition interaction: *F*(1,52) = .22, *p = .644*.

**Supplementary Figure captions**

*Supplementary Figure 1.* Distributions and means of RT (panel A), and accuracy (panel B) per condition (congruent and incongruent) in the flanker task for both monolinguals and bilinguals. Error bars represent 95% confidence intervals.
